# Engineering surface electrostatics affords control over morphological preference, synergy, and activity in polymer degrading enzymes

**DOI:** 10.1101/2025.01.06.631575

**Authors:** Liliana Oliveira, Elaine M. Rudge, Michael Zahn, Victoria Bemmer, Kerry R. Green, Andrew R. Pickford, Bruce R. Lichtenstein

## Abstract

The biocatalytic recycling of plastics, such as polyethylene terephthalate (PET), promises a sustainable alternative to our present open-loop cycles. Engineering of PET-hydrolases for this purpose has focused on improving activity near the glass-transition temperature of the polymer by increasing their thermostability, neglecting other features of the protein-polymer system that affect enzymatic activity. Here, we isolate the effect of electrostatics on the activity of a thermophilic PETase by rationally redesigning its surface charge, while preserving its thermodynamic properties. The enzyme variant, *Sf*Inv, shows orders of magnitude improvements in binding affinity and in activity towards untreated plastic films, with inverted morphological preference. When combined, the wildtype enzyme and *Sf*Inv act synergistically, revealing an entirely new mechanism for cooperative activities driven by complimentary electrostatic interactions at the PET surface. These findings highlight unexplored avenues in improving PETase function through the control of morphological preference or introduction of protein cooperativity by exploiting protein electrostatics.

## Introduction

Using enzymes for biocatalytic depolymerisation of plastics provides a low-energy and low-resource alternative to the current open-loop recycling of common consumer materials^1^. In the case of polyethylene terephthalate (PET), this process allows for the nearly quantitative recovery of the constituent monomers terephthalic acid (TPA) and ethylene glycol (EG), making new routes to (bio)-chemical valorisation and repolymerisation accessible^2,3^. The range of PETases, enzymes capable of carrying out this conversion, has also been expanded in recent years through discovery and further engineering^4–8^. Most of these research efforts have targeted large-scale bio-recycling through increases in enzymatic thermostability, allowing for improved reaction rates and depolymerisation extents by taking advantage of higher polymer chain mobility near the bulk PET glass transition temperature (T_g_)^9–14^. This primary focus on thermostability has deepened our understanding of thermodynamically important structural features of PET-degrading enzymes;^10,14^ however, it has left obscure much of the fundamental nature of the interactions between the proteins and the polymer surface, which ultimately governs enzymatic selectivity, optimal conditions and industrial performance.

Plastic-degrading enzymes face a multitude of challenges associated with the physical-chemical properties of their solid, hydrophobic substrate beyond polymer chain dynamics, which cannot be overcome by enhanced enzymatic thermostability and higher reaction temperatures. On solid substrates, catalytic activities can be influenced by restricted diffusion, variations in the density of surface charge, degree of crystallinity, and protein-protein interactions^15–19^. These complications have been shown to underlie some of the more counter intuitive behaviours we observe in natural PETases, such as diminished activities at elevated enzyme concentrations,^17,20,21^ which prevent some members of the enzyme family from being used at an industrial scale. Studying such attributes can therefore clarify fundamental enzyme-polymer interactions underlying catalytic activities, allowing us to better engineer biocatalysts towards industrial recycling applications.

In this study we examine *Sf*Cut, a highly active, thermostable PETase from *Saccharopolyspora flava* most closely related to PHL7^8^ (Figure 1), previously reported^4^ to have an exceptional preference for micronized PET powder over amorphous PET film. We sought to understand the molecular basis for this morphological preference, hoping to gain insight into how substrate specificity is controlled. Through focused structural analysis, we identified that the principal feature differentiating *Sf*Cut from other PETases was its markedly negative charge; to explore the implications of this property, we used a rational protein design approach that maintained the thermodynamic stability of *Sf*Cut while inverting its overall charge. This allowed us to clarify the role of protein electrostatics in substrate morphological preference and provided new insights into how we can use protein charge to exploit protein-protein interactions at the plastic surface. By connecting structural analysis with careful protein design, this study contributes to advancing the development of more efficient targeted strategies for improving enzymes for plastic biodegradation.

**Figure 1.**
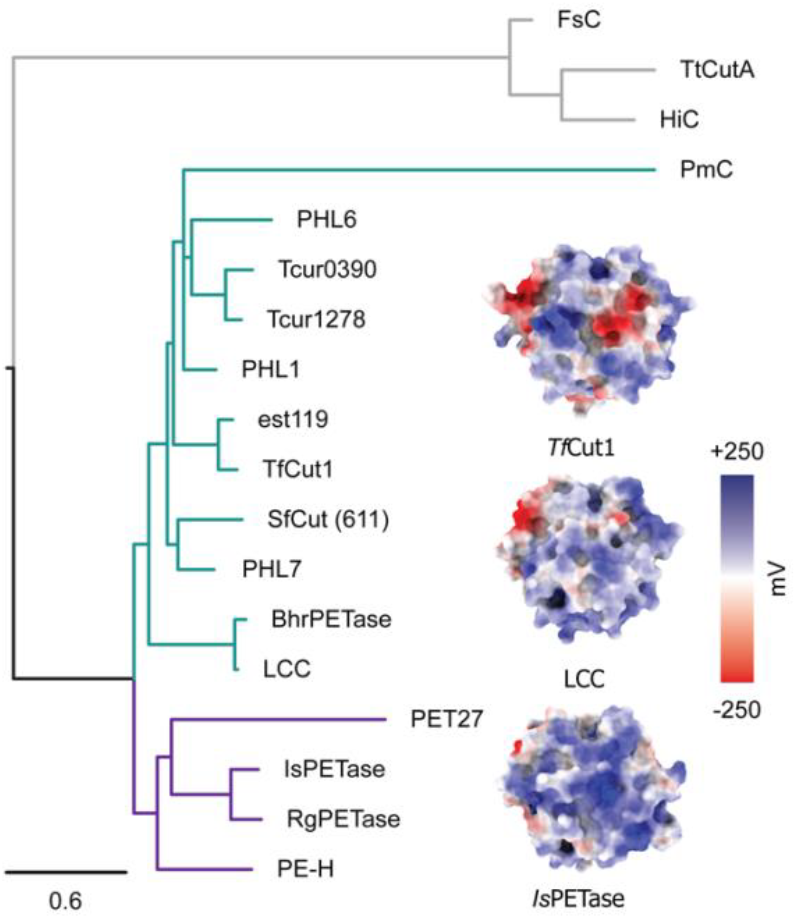
Phylogenetic tree showing relationship between *Sf*Cut and other previously reported PETases. The structure of three well characterised PETases are represented as surfaces coloured by their surface potential at pH 7.5, as calculated by APBS^*51*^. *Sf*Cut falls within the same clade as the thermotolerant type I PETases (green). Type II bacterial PETases are shown in purple and those of fungal origin are shown as the grey outgroup. Accession codes for the PETases used can be found in Supplementary Table 1 and the alignment used to build the tree in Supplementary Figure 1.

## Results and Discussion

### Structural analysis of *Sf*Cut and surface charge redesign

To understand the parameters that influence the substrate morphological preference of *Sf*Cut, we carefully examined its structure. Like other reported PETases, *Sf*Cut (PDB: 7QJP) adopts a typical alpha-beta hydrolase fold, with a conserved disulfide bond between Cys243 and Cys260. The active site matches that of a typical PET-degrading serine hydrolase, composed of a catalytic triad (Ser132, Asp178 and His210), a conserved tryptophan (Trp157), an oxyanion hole (Phe64 and Met133) and a lipase box. Despite these features, *Sf*Cut is strikingly different to other highly active PETases owing to its negative surface charge and total charge density of -0.55 per kDa at neutral pH. These charges are mostly found on the surface opposite to the active site, away from residues expected to directly interact with the plastic (Figure 2a/c). This pronounced charge appears to modulate the morphological selectivity of *Sf*Cut, causing its activity to drop considerably between pH 6.0 and pH 7.5 on films while it increases on amorphous powders^4^. As PET has a stable, negative zeta potential independent of pH at these conditions^22^, the changes in depolymerisation rates implicate the involvement of both protein and polymer electrostatics as significant factors in these properties.

**Figure 2.**
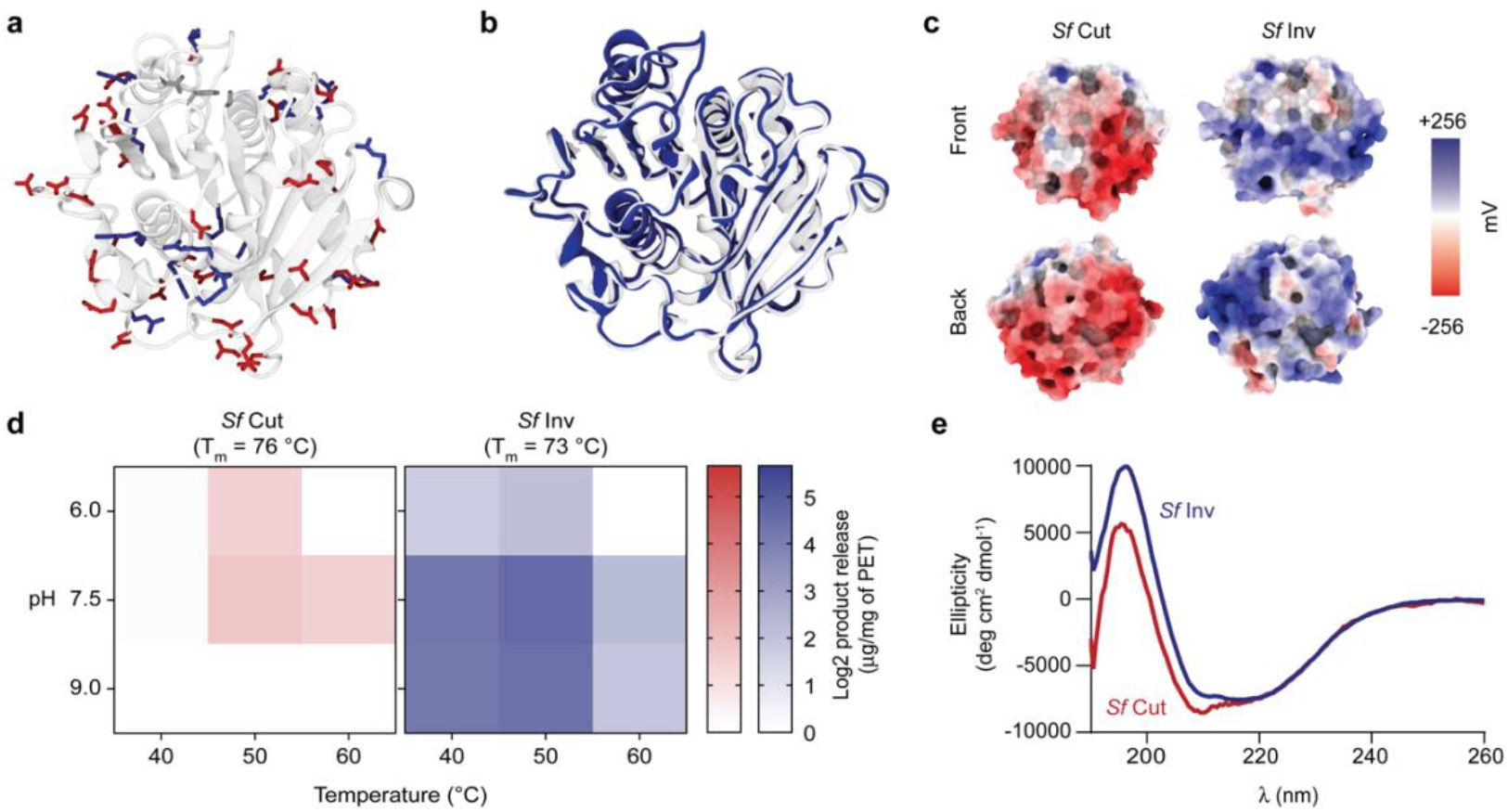
Biophysical characterisation of *Sf*Cut and *Sf*Inv. **a** Cartoon representation of *Sf*Cut, with positively and negatively charged residues represented as sticks. Positive charges are shown as blue, negative charges as red. Active site residues are shown in grey at the top of the structure. **b** Structural alignment between *Sf*Cut (white, PDB: 7QJP) and the crystal structure of *Sf*Inv (blue, PDB: 9EWR) confirms the adoption of an alpha-beta hydrolase fold and the success of the design with a Cα RMSD of 0.36 and a TM-score of 0.99. **c** Surface representation of the potential surfaces of *Sf*Cut and *Sf*Inv at pH 7.5, as calculated by APBS. **d** Heatmap for activity optima of *Sf*Cut and *Sf*Inv at 100 nM on amorphous PET powders across temperatures and pH. Apparent T_m_ at a scan rate of 90 °C/hour for the respective enzymes is indicated. Additional data for PET films and semi-crystalline powders can be found in Supplementary Figure 7. **e** Circular dichroism spectra at 50 °C shows both enzymes remain well structured at their reaction optima.

We aimed to investigate how protein charge influences substrate morphology preference of *Sf*Cut by extensively redesigning its surface. To ensure clarity in our analysis, we sought to isolate the effects of charge substitutions from inadvertent changes to enzyme activity caused by variations in protein thermostability. To avoid this, we selected a manual design approach over computational methods, as the later have been constructed in a way that produces stabilised proteins as outputs. For design purposes, all charged residues in the crystallographic structure, including histidines involved in salt bridges, were identified and counted towards the overall protein charge (-16). Owing to the lack of observed crystallographic density, we did not count the first and last glutamic acid residues in the sequence.

To minimise disruption of the protein-polymer interface and the active site, only residues at a distance greater than 10 Å from the catalytic triad were considered for mutation. For the purposes of design, we ranked residues based on how amenable to change they might be, excluding those that were buried, involved in critical polar contacts, stabilising secondary structural elements, or providing an ambiguous structural role. Our choices of sites for mutation were supported by referencing the position-specific scoring matrix (PSSM) of *Sf*Cut (Supplementary Spreadsheet 1) excluding residues at highly conserved sites. As a priority for mutation, we focused on changing isolated negatively charged residues on *Sf* Cut’s surface, considering structural context to minimise steric clashes and preserve native contacts. Once all the free residues amenable to change were exhausted, we proceeded to consider mutations within larger salt-bridge clusters followed by neutral surface residues. In the interest of investigating solely the effect of the surface charge, we did not intentionally introduce mutations that could create new stabilising interactions.

The redesigned variant, here referred to as *Sf* Inverse (*Sf*Inv), incorporates 24 mutations (Supplementary Table 2; Supplementary Figure 2) at sites with an average information content of 0.41, resulting in an overall charge of +16 (Δ_charge_ of +32) and a charge density of + 0.54 per kDa at neutral pH. Structural predictions using ColabFold^23,24^ and ESMfold^25^ (pLDDT > 0.9, Supplementary Figure 3) closely align with the crystal structure of *Sf*Cut, having C_α_ RMSD values of 0.38 Å and 0.52 Å, respectively. These results suggested that the redesign was likely to preserve the protein fold, without introducing structural changes that could impact enzymatic function.

### Biophysical characterisation of *Sf*Inv

We sought to confirm that the extensive surface modifications introduced to *Sf*Inv did not impact its biophysical properties or its ability to hydrolyse PET plastic. *Sf*Inv expressed in *E. coli* at comparable levels to the wildtype enzyme, and its identity was confirmed by mass spectrometry after purification (Supplementary Spreadsheet 2). Both *Sf*Inv and *Sf*Cut showed similar melting temperatures and thermo-kinetic profiles when analysed by DSC. The apparent T_m_ were 73 °C for *Sf*Inv and 76 °C for *Sf*Cut (Figure 2d, Supplementary Figure 4, Supplementary Table 3), with both enzymes unfolding in a single irreversible step therefore realising our design ambition of minimising the impact on thermodynamic stability.

The protein fold and success of the design were further confirmed through circular dichroism (CD) and structure determination by X-ray crystallography. *Sf*Inv readily crystallised under several conditions, and its structure was solved to a final resolution of 1.17 Å (Supplementary Table 4). The asymmetric unit in the structure contained two chains, which aligned with a C_α_ RMSD 0.24 Å, and the more complete chain B was used for structural analysis. The solved crystallographic structure of *Sf*Inv (PDB: 9EWR) confirmed that the enzyme retained its alpha-beta hydrolase fold (Figure 2b) and exhibited the expected positively charged surface (Figure 2c). Structural alignment with the wildtype protein yielded a C_α_ RMSD 0.36 Å and a TM-score^26^ of 0.99 (Figure 2b), demonstrating that the overall fold and structure were preserved, despite extensive modification. We were able to confirm that no new stabilising interactions were introduced, and that the backbone conformation, disulfide bond, and salt-bridges remained largely intact, with the only exceptions being due to crystallographic contacts spanning symmetry related monomers (Supplementary Figure 5). CD also revealed that both proteins remain well folded under the reported optimum reaction temperature of *Sf*Cut (50 °C) (Figure 2e).

With confidence in the achievement of our structural design goals, we explored the activity of *Sf*Inv on amorphous PET powder. *Sf*Inv exhibited an optimum reaction temperature at 50 °C, similar to that of the wildtype enzyme (Figure 2d), but demonstrated a higher pH optimum (pH 9) compared to *Sf*Cut (pH 7.5). This shift to a higher pH optimum aligns with the activity profiles of positively charged PETases, which demonstrate pH optima above pH 8^4,27^. We anticipated that the changes in the surface charge between *Sf*Cut and *Sf*Inv could lead to substantial differences in their binding affinity to PET. Indeed, despite its relatively high activity, *Sf*Cut showed no appreciable binding to the amorphous PET powder under tested conditions (Figure 3b, Supplementary Figure 6). This stands in clear contrast to *Sf*Inv, which demonstrated a K_d_ of less than 4.5 nM with a surface coverage (Γ_max_) of 10-16.5 nmol g^-1^ PET, consistent with characteristics measured with other highly active PETases^28^. These results confirmed the success of our design efforts, and provided a basis for us to examine the isolated effects of protein charge on PET-degrading activities and morphological preference.

**Figure 3.**
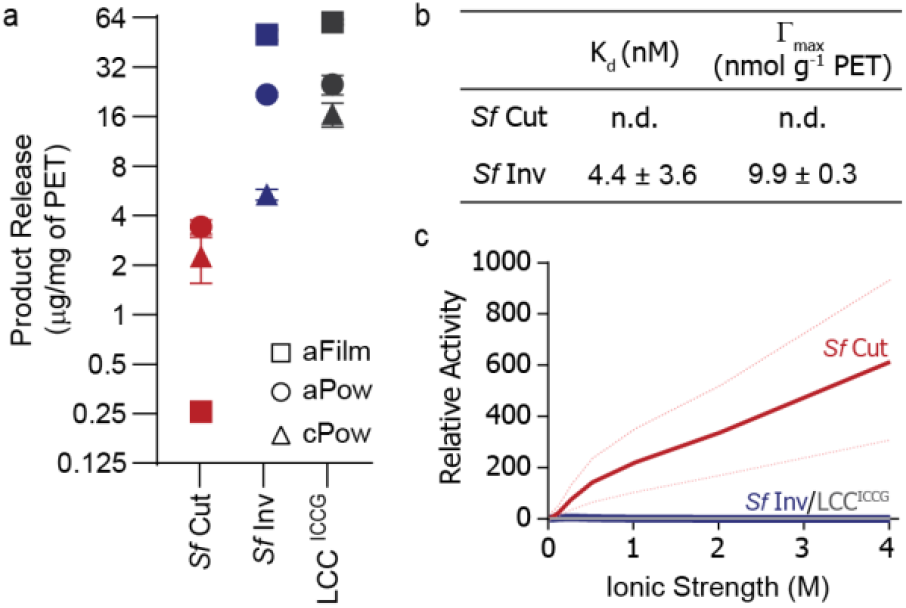
Investigating how ionic strength plays a role in protein-polymer interactions. **a** Substrate preference for *Sf*Cut, *Sf*Inv, and LCC^ICCG^ on a Log_2_ scale. Experiments were carried out at the optimum temperature and pH for the individual enzymes at 100 nM, with 100 mM sodium chloride. **b** *Sf*Inv has a measurable affinity to PET powders, while *Sf*Cut interacts too weakly to measure (binding curves and full statistics are detailed in Supplementary Figure 4). **c** Ionic strength affects the relative activities of *Sf*Cut and *Sf*Inv on aFilms in different ways: *Sf*Cut benefits from the increased ionic strength, whilst *Sf*Inv, like LCC^ICCG^, is largely not affected. Plot is shown as relative activity with respect to the activity of each enzyme in buffer without added salt. Lighter dotted lines represent standard deviations calculated with propagated error.

### The role of surface charge on protein-polymer interactions

To explore how their significant difference in protein surface charge manifested in terms of depolymerase activity across PET morphologies, we examined *Sf*Cut and *Sf*Inv on three PET substrates: amorphous powder (aPow), amorphous film (aFilm) and semi-crystalline powder (cPow) (Figure 3a, Supplementary Figure 7). *Sf*Inv demonstrated higher activity than the wildtype enzyme across all substrates tested, with broader pH and temperature optima, as well as three-fold and six-fold more product release on semi-crystalline powders and amorphous powders respectively. Our results confirmed previous observations that *Sf*Cut has a considerably lower activity on films compared to powders, independent of substrate crystallinity. In contrast, *Sf*Inv showed an inverted selectivity: its activity on films was enhanced, approximately 2-fold higher than on powders, demonstrating that enzyme specificity for different substrate morphologies is a genetically encoded, engineerable property. This corresponds to over 200-fold enhancement in the activity on amorphous PET films compared to *Sf*Cut, effected simply by surface charge inversion. Despite the substantial difference in evident binding affinity to amorphous powders, both enzymes show activity saturation on amorphous films above 250 nM, confirming that binding alone is insufficient to explain their catalytic properties (Supplementary Figure 8). Although not an intended outcome of our design process, *Sf*Inv demonstrated a similar level of conversion as the most promising industrialised enzyme, LCC^ICCG, 10^, at their respective optima (Figure 3a). In a direct comparison at 50 °C, *Sf*Inv outperformed LCC^ICCG^ at laboratory scale (Supplementary Figure 9).

Although the greatly enhanced binding affinity observed with *Sf*Inv can partially explain its higher enzymatic activity, differences in enzymatic morphological preference must be caused by a change in how the differently charged enzymes interact with the polymer. Electrostatic interactions between the polymer surface and the enzymes can be mediated by both unspecific and specific interactions with ions in solution, therefore we assessed the activities of both enzymes at a range of ionic strengths, using both monovalent and divalent salts. The negatively charged wildtype enzyme, *Sf*Cut, showed a monotonic rise in activity on amorphous films as ionic strength increased from 0 M to 4 M, with a relative activity enhancement of nearly 600-fold (Figure 3c, & Supplementary Figure 10a). This behaviour contrasts with that of some of the best-performing PETases, such as LCC^ICCG^, where the depolymerase activity is not affected by the ionic strength of the solution (Figure 3c & Supplementary Figure 10b). For *Sf*Inv the effect of ionic strength was less pronounced (Figure 3c), with activity peaking at 250 mM ionic strength at approximately five-fold higher than in buffer with 0 M salt, before declining to approximately 7% of its maximal value at 2 M (Supplementary Figure 10a). These results on amorphous films were consistent across different salts (Supplementary Figure 10), indicating that the selectivity observed in *Sf*Cut and *Sf*Inv is driven by electrostatic interactions between the enzyme and the polymer surface, rather than by specific interactions mediated by salts.

Interestingly, the effect of ionic strength on the enzyme activity was distinct on amorphous powders. *Sf*Inv showed the same monotonic rise in activity as *Sf*Cut, although less pronounced (Supplementary Figure 10c), with both enzymes seeing no benefit to activity above an ionic strength of 1 M. Overall, positively charged *Sf*Inv sees relatively moderate benefits of increased ionic strength, with a greater effect observed on powders than films, whereas *Sf*Cut shows substantial improvements in activity with salt, benefitting more on films than powders. This demonstrates that the interplay between the electrostatic fields of the polymer and the protein varies between polymer morphologies, influencing the observed selectivity of the two enzymes, with notably more dominant effects on PET film digestions.

Despite the benefit of ionic strength on the activity of *Sf*Cut, it is important to note that the total product released by *Sf*Cut did not surpass that of *Sf*Inv at its maximum under the conditions tested. Additionally, while increased salt concentration did have an effect on the apparent T_m_ of *Sf*Cut, causing an increase of 4 °C at 2 M sodium chloride, no such change was observed for *Sf*Inv (Supplementary Figure 11). This suggests that some of the activity improvement in *Sf*Cut with increasing ionic strength may be a result of enhanced thermostability, an effect that is absent in *Sf*Inv.

### *Sf*Inv shows greatly enhanced PET-degrading activity at pilot scale and on post-consumer waste

The improved selectivity of *Sf*Inv towards amorphous PET films under analytical conditions suggested a potential for digesting unmodified amorphous films at pilot scale, pH-controlled experiments. Under these conditions (1 mg_enzyme_ g^PET-1^ with high solids loading of 20% w/v)^1,29^, the wildtype enzyme (*Sf*Cut) demonstrates limited activity, achieving less than 1% substrate conversion and a monomeric product yield of 0.15 g/L as measured by HPLC. In contrast, the engineered *Sf*Inv exhibited significantly higher efficiency, achieving approximately 8% substrate conversion within 24 hours (Figure 4a), with a monomer yield of 18.6 g/L. Strikingly, at substantially lower enzyme loading (1 µM, ∼0.15 mg_enzyme_ g^PET-1^), *Sf*Inv achieved the same high levels of conversion within the same time frame, suggesting there is substantial potential for reaction condition optimisation to maximise the digestion of amorphous PET films while reducing resource requirements (Supplementary Figure 12).

**Figure 4.**
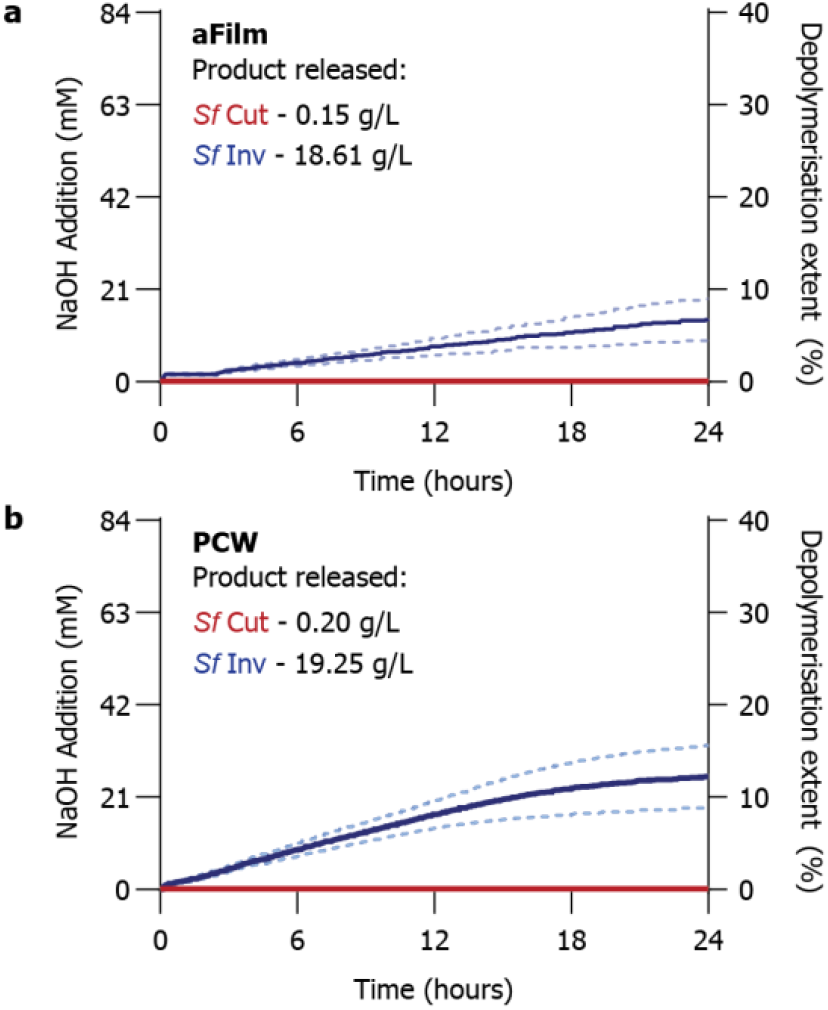
Depolymerisation of PET at pilot scale by *Sf*Cut and *Sf*Inv. Enzymes were incubated with 20% (w/v) PET substrate, at an enzyme loading of 1 mg per g of PET. *Sf*Cut is shown in red, whereas *Sf*Inv is shown in blue. Figure illustrates depolymerisation overtime of amorphous PET film (**a**) and post-consumer PET waste (**b**). Average trace showed as a solid line, each of the replicates are represented as dotted lines in the corresponding colour. Product release as determined by HPLC shown in each panel.

Comparable results were also found to be true on untreated post-consumer waste (PCW) in the form of PET-film sandwich trays (Figure 4b, Supplementary Table 5, Supplementary Figures 13 and 14). *Sf*Inv efficiently digested over 10% of the waste within 24 hours, achieving a measured yield of 19.25 g/L, while *Sf*Cut yielded negligible amounts of product (Figure 4b). This further highlights that it may be possible to engineer enzymes specifically targeting the processing of complex post-consumer waste streams of PET with limited pre-treatments, upstream of current methods reliant on resource-intensive preparation of micronized powders.^29^

### *Sf*Cut and *Sf*Inv act synergistically in degrading aPET films

Building on the evident enhancement in activity and change in selectivity of *Sf*Inv over *Sf*Cut on PET substrates, we sought to understand whether the enzymes act at distinct sites on the PET surfaces with differing geometries or electrostatic charge, which could account for the observed differences in morphological preferences and allow for the enzymes to work synergistically in degrading PET. Specifically, we hypothesised that the amount of product released from a mixture of the two enzymes could exceed the sum of the product release for each enzyme individually at its respective concentration. To quantify this, we define synergy as:

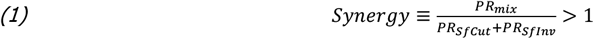

where PR_*Sf*Cut_, PR_*Sf*Inv_, and PR_mix_ are the product released by each enzyme at specified concentrations, and their mixture, respectively. When *Sf*Cut and *Sf*Inv were mixed at different concentrations and ratios, and applied to PET films, we observed clear evidence of synergistic activity (Figure 5a). The effect is most notable at lower concentrations of *Sf*Inv, with a synergy value above 2 (Figure 5c). However, at elevated concentrations of *Sf*Inv, the synergistic benefit is lost, possibly owing to surface crowding^17,20^ or competition for binding sites on the plastic substrate. In contrast, increasing *Sf*Cut does not eliminate the observed synergy, suggesting that each enzyme plays a distinct role when acting in concert on the film surface.

**Figure 5.**
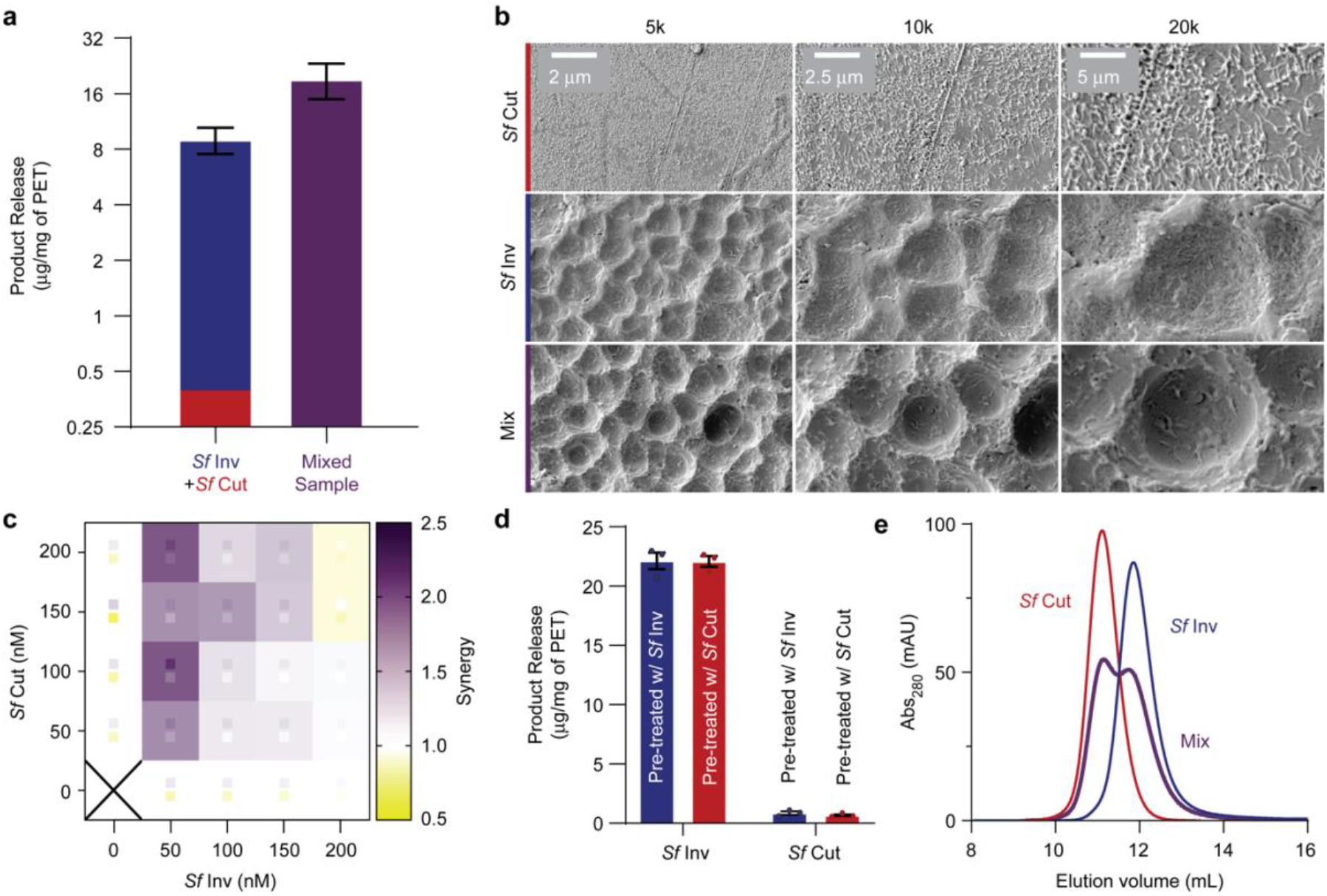
Investigating the synergy between *Sf*Cut and *Sf*Inv on aPET film. **a** Difference in total product release by pure enzymes and their mixture, at the condition with highest synergy (100 nM:50 nM of *Sf*Cut to *Sf*Inv). Pure enzymes are represented as a stacked plot, where red represents *Sf*Cut and blue *Sf*Inv. The enzyme mixture is represented as a purple bar. Product release is shown as a Log_2_ scale for clarity. Error bars represent 1 SD. **b** Scanning Electron Microscopy (SEM) images of amorphous PET film after incubation with enzyme. The top two panels show the film after incubation with *Sf*Cut or *Sf*Inv individually, whilst the bottom panel shows the film after incubation with the enzyme mix. **c** Heatmap of the synergy ratios across different concentrations of *Sf*Cut and *Sf*Inv, with the purple as productive synergy and yellow as a decrease in total product release. Standard deviation is shown as smaller squares in the same colour scheme. **d** Total product release for the stepwise synergy, where the pure enzymes were incubated with an amorphous film coupon pre-digested by either *Sf*Inv (blue) or *Sf*Cut (red). Error bars represent 1 SD. **e** Chromatogram from Size Exclusion Chromatography (SEC) for *Sf*Cut (red), *Sf*Inv (blue) and a mixed sample of both enzymes at equimolar ratios (purple).

To further clarify the mechanism of the synergistic activity, we explored potential causes involving the differing enzyme charges. One possibility was that the enzymes might assemble in solution in a way that enhances their stability and activity. To test this, we incubated the enzymes together at elevated concentrations (∼ 34 µM, compared to the 0.05-1 µM enzyme concentrations used in polymer digestions) and evaluated whether they form complexes by size exclusion chromatography on the mixture (Figure 5e); no shift was observed in the elution profile, suggesting that the enzymes do not appreciably bind to each other in solution. To exclude the possibility that the enzymes are targeting different sites on the polymer chain and therefore modifying the plastic surface such that it becomes a better substrate for the oppositely charged enzyme, we examined whether the enzymes showed a stepwise synergy. The activities of the enzymes were evaluated on amorphous PET film coupons that had been pre-treated with either *Sf*Cut or *Sf*Inv (Figure 5d), with no significant difference in activity observed on films pre-treated with either enzyme, showing that the surface targets for both enzymes are largely the same.

Given these findings, and that *Sf*Cut alone shows negligible binding to PET, it is likely that the synergy observed at principally low concentrations of *Sf*Inv is a result of charge masking, similar to that of the ionic strength experiments. In this case, *Sf*Inv binds to the plastic surface masking the negative charges of the polymer and any bound *Sf*Cut, which in turn allows more *Sf*Cut to bind productively. Similar effects have been previously observed with the use of synthetic surfactants^30^, where anionic detergents were used to increase the negative surface charge of the polymer, therefore attracting the cationic *Is*PETase^31^ to the surface, accelerating enzymatic degradation. In this case, however, the enzymes themselves are modulating the surface charge.

The difference in how the film surface is modified by the enzymes acting individually or as a mixture can be visualised after partial digestion of PET coupons using SEM (Figure 5b). *Sf*Cut increases the roughness of the surface leaving what appears to be thread-like crystalline regions behind, whilst *Sf*Inv shows broad pitting, a surface modification commonly observed with other highly active PETases^18,31^. However, with the enzyme mixture, an unusual double-pitting effect is observed. Broad pits similar in dimensions to those formed by *Sf*Inv contain deeper pits within them, perhaps from *Sf*Cut being ‘funnelled’ into the centre of areas where the positively charged *Sf*Inv masks the negative charge on the surface of the film.

As far as we are aware, this is the first time that synergy has been observed in enzymes directly modifying plastic surfaces. These results reveal a previously unexplored mechanism by which enzyme mixtures can enhance the degradation of plastic through beneficial electrostatic interactions, offering new opportunities for improving bio-recycling through synergistic enzymatic activity.

## Conclusions

Prior to this study the most significant changes in the activity of PET degrading enzymes were realised by increasing the performance of the biocatalysts at elevated temperatures, where the polymer is more dynamic. This focus on thermostability has obfuscated how other physical-chemical properties of the polymer surface influence the activity of PETases, limiting our understanding on the enzyme-substrate interactions at play.

Here, by rational redesign of the surface of a highly active PETase, *Sf*Cut, we were able to investigate the impact of electrostatics on enzymatic activity in the absence of confounding factors like thermodynamic stability or changes in features near the active site^13,30,32^. Through carefully crafted studies on the effects of ionic strength on product released from different substrates, we demonstrate that substrate selectivity is driven by polymer morphologies having distinct electrostatic profiles, and therefore influencing enzyme-surface interactions in contrasting manners. Measured zeta potentials are known to have smaller magnitudes on rough surfaces than on smooth surfaces^33,34^, and the impacts of this are likely manifesting here. Within the diffuse layer around the neatively charged polymer surface, the anionic *Sf*Cut experiences repulsion while the cationic *Sf*Inv experiences attraction, with these interactions being more pronounced on smooth films than on rough powders.

By focusing on electrostatic interactions governing the activity of PET degrading enzymes at the polymer surface, we demonstrated that it is possible to not only tune the selectivity of PETases for substrates of differing morphologies, but also increase their binding affinity through the rational engineering of surface charge away from the active site. The process of introducing large scale changes in protein surface electrostatic potential in *Sf*Inv also afforded substantially improved activities on all PET substrates tested, exposing the benefits of using an electrostatically-oriented approach to engineering plastic depolymerases for improved activities at the low salt concentrations relevant for industrial processes. As we demonstrate through the preservation of the thermodynamic stability of *Sf*Inv, this approach is complementary to established methods designed to enhance the thermostability of enzymes and can be introduced in the engineering process towards better industrial PET depolymerases.

We also found that synergistic activity between PETases can be realised through the applications of complementarily charged proteins. While the enhancement in activity we observed on films is relatively modest, just over two-fold, this advance is the first method for building synergy into enzymes acting at plastic surfaces and opens the door to engineering campaigns specifically tailored towards improving it. In contrast to synergies between enzyme pairs acting at different chemical sites on a substrate, the mechanism of synergy in this case appears to depend upon the electrostatic masking effect of the positively charged *Sf*Inv when bound to PET, which in turn promotes the activity of the negatively charged *Sf*Cut. This mechanism is likely generalisable and opens the possibility of exploiting this discovery to reduce the enzyme loading needed for industrial processes or to create dynamic mixtures of enzymes capable of accommodating variations in surface electrostatics across substrates or over the course of depolymerisations.

Through isolating the effect of electrostatics on *Sf* Cut’s activity on PET, we have been able to establish an engineering approach that allows tuning of binding affinity, improvement of enzymatic turnover, control of substrate morphological preference, and introduction of functional synergy in PETases. These insights not only expand the repertoire of established features under the control of rational protein design for plastic depolymerases, but are also likely not limited to PET. As such, we expect these results to be translatable to enzymes capable of digesting a broad range of synthetic polymers, and that the considerations established here will prove crucial when finding and engineering enzymes to tackle plastics found in complex industrial and post-consumer waste streams.

## Materials and Methods

Amorphous PET film (ES30-FM-000145) and semi-crystalline PET powder (ES30-PD-006031) were purchased from Goodfellow. Post-consumer plastic waste was obtained from PET sandwich packaging. All reagents for molecular biology and strains were purchased from New England Biolabs. All other reagents and buffer components were acquired from Fisher Scientific or Merck, unless stated otherwise.

### Phylogenetic and sequence analysis

Protein sequences of reported PETases were aligned using ClustalOmega^35^ with default settings. A phylogenetic tree was built from the sequence alignment using IQtree^36^ with 100,000 UltraFast bootstraps^37^, nearest neighbour interchange (NNI) search, automatic model selection^38^, and 100,000 cycles of single branch testing (SH-aLRT)^39^. Sequence identities were found using protein BLAST^40^ with default settings. The position specific scoring matrix was calculated in POSSUM^41^ using Uniref50 as the database, with 3 iterations and an E-value threshold of 0.001.

### Manual surface redesign

The surface redesign was done manually using the crystal structure of *Sf*Cut (611, PDB: 7QJP). All charged residues with crystallographic densities were identified, including histidines involved in salt bridges or catalytic contacts; those that were not interacting were counted as non-charged. Redesign focused on charged surface residues more than 10 Å away from the active site not involved in any polar or evident structural contacts, when these were exhausted mutations within larger salt-bridges and to uninvolved surface neutral residues were considered. No mutations were added to purposefully increase or impair enzymatic stability and activity. The overall charge was changed from -16 to +16, by mutating negatively charged as well as neutral surface residues mostly not involved in stabilisation of the protein structure by visual inspection. The final *Sf*Inv sequence was modelled using ColabFold/AlphaFold2^23,24^ and ESMFold^25^ to confirm that no disruption to the protein structure by the design was predicted.

### Plasmid construction

Genes for *Sf*Cut and *Sf*Inv were synthesised by Twist Bioscience. *Sf*Cut was cloned by Twist directly into pET21b(+), *Sf*Inv was synthesised as a gene fragment and cloned into pET28b(+) using Gibson assembly. The assembly mixture was transformed into NEB5α competent cells, DNA purified (Qiagen miniprep kit), and sequence confirmed by Sanger sequencing (Eurofins Genomics). Both constructs include a C-terminal His-tag, and were sequence optimised for *Escherichia coli*.

### Protein expression and purification

Proteins were expressed using BL21(DE3) *E. coli* strain. Cells were grown in terrific broth with the selection antibiotic at 37 °C, in 4.5 L cultures in bioreactors (Eppendorf BioFlo 120w) with pH and air flow control (Biocommand Bioprocessing Software). Protein expression was induced at an OD_600_ of 1.2 for 18 hours at 20 °C, using a final concentration of 1 mM IPTG. Harvested cells were resuspended in HisTrap binding buffer (20 mM Tris-HCl pH 8.0, 300 mM sodium chloride, 40 mM imidazole) with nuclease (expressed in house) and 25mM of magnesium added. The resuspended cells were then homogenised, sonicated (Amplitude 40, 3 sec ON, 9 sec OFF for a total processing time of 6-10mins), and clarified by centrifugation at 55,000 x g. Clarified lysate was filtered through a 0.45 µm MCE filter and purified by affinity chromatography on a HisTrap FF (5 mL) column, eluted over a gradient up to 500 mM imidazole. The protein peak was further purified by size exclusion chromatography using a Superdex 16/600 HiLoad 75pg equilibrated with 50 mM sodium phosphate pH 7.5, and 100 mM sodium chloride. SDS-PAGE was run to assess purity.

### Differential scanning calorimetry

Apparent melting temperature (T_m_) values for the purified proteins were determined using a MicroCal PEAQ-DSC with automated sampler (Malvern Panalytical), using a buffer matched to that of the size exclusion chromatography step as reference. The analyses were performed using 1 mg/mL of protein, at a temperature range of 30-100 °C, using low feedback, at 192 °C/hour, 96 °C/hour, 90 °C/hour, 48 °C/hour, 24 °C/hour, 12 °C/hour and 6 °C/hour. Baseline subtraction was performed using the instrument’s data analysis software. Calfitter 2.0^42^ was used to derive the activation energy (E_act_), the heat capacity change (ΔC_p_), the activation enthalpy change (ΔH‡), and the reference temperature of the irreversible melting step (T_act_) using the thermal denaturation model with the lowest SSR value.

### Structure determination by X-ray crystallography

*Sf*Inv was concentrated to 10 mg/mL and crystallised by the sitting drop vapour diffusion method using a Mosquito crystallisation robot (SPT Labtech) and SWISSCI 3-lens low profile plates in condition C8 of the SaltRx screen (Hampton Research): 0.1 M Tris pH 8.5 and 3.5 M sodium formate. Crystals were cryo-protected with 20% glycerol before flash-freezing in liquid nitrogen. Diffraction data were collected at the Diamond Light Source (Didcot, UK) at beamline I03 and automatically processed with the AutoPROC+STARANISO^43,44^ pipeline on ISPyB. The structure was solved by molecular replacement on CCP4 Cloud using Molrep^45^ and an AlphaFold2-model^23^. Coot was used for model building, followed by model refinement using Refmac^46^. The final structure was evaluated with MolProbity^47^, and the structure has been visualised in VMD^48^ and ChimeraX^49^. The structure was deposited in the PDB with code 9EWR. Data and refinement statistics can be found in Supplementary Table 4.

### Structural characterisation by circular dichroism

Spectra were collected on a PiStar-180 (Applied Photophysics) with water bath temperature control. Protein samples were analysed at a concentration of 0.1 mg/mL in 10 mM sodium phosphate pH 7.5 with 20 mM sodium chloride at 50 °C, in a 1 mm stoppered quartz cuvette. Data collected at wavelengths between 200 and 260 nm, with half bandwidth of 1.5 nm and a wavelength interval of 0.5 nm (1 sec per point, 5 repeats), were averaged and baseline subtracted using a matched buffer blank. Raw ellipticity data was converted to mean residue ellipticity by dividing by the path length, concentration, molecular weight and number of residues.

### Amorphous PET powder production and analysis

Sheets of amorphous PET film were cut into strips, immersed in liquid nitrogen and cryo-milled at 2,400 rpm in a SM300 cutting mill (Retsch), with a bottom sieve with 4 mm square holes. Subsequently, this product was reduced in size further by immersing in liquid nitrogen and cryo-milling at 18,000 rpm in a ZM200 centrifugal mill, with a 0.12 mm ring sieve with trapezoidal holes. The particle size and crystallinity of the cryo-milled amorphous PET powder was compared to that of the purchased semi-crystalline PET powder, as well as the PET film, using a CAMSIZER X2 (Microtrac MRB) and Differential Scanning Calorimetry (Supplementary Table 5, Supplementary Figures 13 and 15).

### Reaction quenching and product quantification by HPLC

All reactions were quenched by addition of equal volume of HPLC-grade methanol, and PET solids removed. Samples were centrifuged at 10,000 x g, using a table-top centrifuge, prior to analyte quantification by HPLC. For samples with subsequent polymer analysis, the PET was washed by rinsing three times with a 1% (w/v) SDS solution, followed by multiple rinses with distilled water. The partially digested polymer substrates were then dried at room temperature under vacuum before analysis. The HPLC analysis was adapted from a reported UPLC method^14^ to allow for HPLC pressures as previously described^50^. Samples were evaluated on a pre-equilibrated C18 Kinetex LC column (00B-4605-AN) with a guard, at 1.1 mL min^-1^ with 0.1% formic acid and acetonitrile as the stationary and mobile phases respectively. Samples were prepared with a known dilution to an absorbance at 240 nm of around 1.0 before 10 µL were loaded onto the column using an automatic sampler (Agilent). Samples were eluted with an isocratic elution at 13% mobile phase for 0.87 minutes, followed by a step to 95% mobile phase for 1.12 minutes and a re-equilibration at 13% mobile phase until a total time of 3.6 minutes. Peaks were integrated using Agilent’s OpenLab software and the product quantification was performed against calibration curves of known standards (TPA, MHET, BHET). An example of the HPLC trace and elution times is provided in Supplementary Figure 16.

### PET degradation assays at small scale

Unless stated otherwise, small scale assays were set up in 1.5 mL tubes with 11 mg of PET substrate, and incubated in triplicate 500 µL reactions for 24 hours at 300 rpm on thermomixers. A final concentration of 100 nM enzyme and 100 mM sodium chloride was used, in 50 mM sodium phosphate pH 7.5, or 50 mM glycine pH 9.0 for *Sf*Cut and *Sf*Inv respectively. In the case of the ionic strength tests 50 mM HEPES pH 7.5 and 50 mM CHES pH 9.0 were used instead, with varying amounts of sodium chloride, magnesium chloride and sodium sulphate. These buffers were chosen in order to minimise their contribution towards the ionic strength. Nevertheless, the buffer component was fully accounted for in the ionic strength calculations by considering their pKas, and calculating the concentration and charges of their respective ionic species. The ionic strength of the solution was calculated by using the formula *I* = ½ *n* ∑_*i*_(*C*_*i*_*Z*_*i*_), where *I* represents the ionic strength, *n* is the number of ions in solution, *C*_*i*_ is the concentration of a specific ion in moles per litre, and *Z*_*i*_ the valence of the particular ion species. For the temperature and pH optima experiments, the reactions were performed at three temperatures (40 °C, 50 °C and 60 °C) and using three different buffers (50 mM MES pH 6.0, 50 mM sodium phosphate pH 7.5, and 50 mM glycine pH 9.0) with 100 mM sodium chloride. To determine the concentration dependency for each of the enzymes, both were tested individually, and together at equimolar ratios, to a final enzyme concentration ranging from 0 - 1 µM.

### Binding isotherms

Binding affinities to the plastic were measure in two ways as described previously^28^. All samples were incubated for 1 hour at 4 °C with rolling in low-binding 1.5 mL microcentrifuge tubes (Eppendorf) to prevent loss from non-specific binding. After centrifugation, free enzyme concentrations were determined using a Micro BCA protein assay kit (Thermo Scientific) with a calibration curve derived for the respective enzyme. 150 µL samples were mixed with the Micro BCA Working Reagent as per the kit’s instructions, and incubated for 1 hour at 50 °C in a covered microtiter plate mixing at 300 rpm. 200 µL samples were measured at 562 nm on a plate reader. *Substrate saturation:* Amorphous PET powder at a fixed substrate loading (100 g/L for *Sf*Cut or 50 g/L for *Sf*Inv) was incubated with 0.05 - 1.5 µM enzyme. The substrate coverage was calculated using the equation 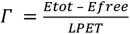, and the K_d_ obtained by fitting the data to the Langmuir adsorption isotherm equation 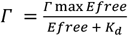, where *Γmax* is the substrate coverage at surface saturation. *Enzyme saturation:* 1 µM enzyme was incubated with substrate loads ranging from 0 – 200 g/L. The bound fraction was calculated from the difference between total and free enzyme concentrations. K_d_ and Γ_max_ were derived from fitting the data to the equation^28^ 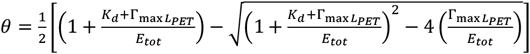, where the fraction bound 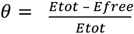, and L_PET_ is the experimental solids loading.

### PET degradation assays – pilot scale and post-consumer waste

Duplicate reactions were undertaken in 250mL MiniBio Reactors (Applikon Biotechnology) to a total reaction volume of 100 mL. Reactions were carried out at 20% (w/v) solids loading and a final enzyme concentration of 1 mg of enzyme per gram of PET. Reactions were incubated at 50 °C, stirring at 200 rpm for 24 hours. Substrates used in the reactions were amorphous PET film or washed and dried post-consumer plastic film waste, roughly cut into 1 cm x 1 cm squares. All pH probes were calibrated immediately prior to analysis, pH changes were followed and maintained by the automatic addition of 1 M (*Sf*Cut) or 5 M (*Sf*Inv) freshly prepared sodium hydroxide via the system pump, which was calibrated prior to the experiment. The base addition was followed as a function of time by the Lucullus^®^ Process Information Management System software and converted to percentage hydrolysis by calculating the moles of TPA neutralised. At the end of the incubation, samples were taken and quenched as described previously using an equal volume of methanol for HPLC analysis. The PET film was recovered by filtration, rinsed thoroughly with deionised water, and allowed to dry before weighing to confirm depolymerisation extent.

### Polymer analysis and characterization

Polymer DSC was performed on a Netzsch DSC 214 Polyma, equipped with aluminium crucibles and lids. Approximately 10 mg of samples was heated from 25 to 300 °C at a rate of 10 °C per minute in a nitrogen atmosphere. Measurements were performed in triplicate, and sample crystallinity was calculated using the equation 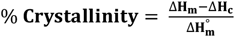, where ΔH_m_ is the enthalpy of melting of the sample ΔH_c_ is the enthalpy of crystallisation of the sample and ΔH_m_° is the enthalpy of melting for a theoretical 100 % crystalline sample (taken as 140.1 Jg^-1^).

### PET degradation assays for investigating synergy

To quantitate synergy across enzyme concentrations and ratios, amorphous PET coupons were incubated in 50 mM sodium phosphate pH 7.5, 100 mM sodium chloride, for 24 hours, with various enzyme ratios and concentrations ranging from 0 – 200 nM, as per Figure 5. To investigate whether the synergy was caused by opposite charge effects between the enzymes, size exclusion chromatography was performed with 1 mg of total enzyme (∼ 34 µM) on its own or at equimolar ratio using a Superdex 75 equilibrated with 50 mM sodium phosphate pH 7.5, and 100 mM sodium chloride. For the stepwise synergy studies, a final concentration of 100 nM enzyme was used. Films were pre-treated with the selected enzyme for 24 hours, reactions quenched and the coupons washed and dried as described above. Coupons were then incubated a further 24 hours with the second selected enzyme before the reactions were quenched one final time prior to product analysis by HPLC.

### Scanning electron microscopy

Samples were incubated with 100 nM of the respective enzyme for 24 hours as described above in PET degradation assays for investigating synergy. For the mixed enzyme samples, a total of 200 nM enzyme at an equimolar ratio was used instead. Samples were quenched, washed and dried as described above before analysis by SEM. PET film samples were mounted onto aluminium stubs using carbon adhesive tabs, and sputter coated with Au/Pd under argon using a Quorum Q150RES (Quorum Technologies Ltd). Samples were imaged using a MIRA3 FEG-SEM Microscope (TESCAN) operated at 3 kV.

## Author contributions

BRL conceptualised and supervised the project. The manuscript was written by LO and BRL, and reviewed by all authors. VB collected polymer DSC data. Enzymes were expressed by EMR. KRG collected crystals for structure determination of *Sf*Inv, and MZ solved the crystal structure. All other data was collected, processed and analysed by LO. ARP and BRL acquired funding.

## Acknowledgements

The authors thank Research England [E3 funding to ARP], the Royal Society [Grant RGS\R2\212336 to BRL], and the UKRI Engineering Biology Mission Hub [Grant BB/Y007972/1 to ARP, BRL and VB]. We thank Diamond Light source for beamtime [PROPOSAL MX-31440], and the staff at beamline I03 for supporting automatic data collection. We thank the Electron Microscopy and Microanalysis Unit at the University of Portsmouth, and in particular Ben Trundle, for assisting with SEM data collection. LO and BRL also thank Prof. Birte Höcker and Prof. Samuel Robson for fruitful discussions and feedback.

