## Supplementary Information for "Engineering surface electrostatics affords control over morphological preference, synergy, and activity in polymer degrading enzymes"

### Table of Contents

#### Supplementary Tables

1. Accession codes for PETases mentioned in this study
2. Amino acid sequences for *Sf* Cut and *Sf* Inv
3. Extended summary of differential scanning calorimetry data with denaturation model fits
4. Data collection and refinement statistics (molecular replacement) for X-ray crystallography structure of *Sf* Inv
5. Summary of polymer differential scanning calorimetry results

#### Supplementary Figures

1. Multiple protein sequence alignment of a set of previously reported PETases.
2. Protein sequence alignment of *Sf* Cut and *Sf* Inv
3. ColabFold/AlphaFold2 and ESM Fold predictions of the *Sf* Inv design
4. Differential scanning calorimetry curves for *Sf* Cut and *Sf* Inv
5. Comparison of polar interactions in modified salt bridges
6. Binding isotherms of *Sf* Inv and *Sf* Cut to amorphous PET powders
7. Heatmaps of optimum pH and temperature for the enzymes across PET substrates
8. Concentration dependence plots for enzymes on amorphous PET film
9. Substrate preference plot for *Sf* Cut, *Sf* Inv and $LCC^{ICCG}$ at 50 °C
10. Ionic strength extended – multiple salts and powder comparison
11. Impact of ionic strength on the apparent $T_m$ of *Sf* Cut and *Sf* Inv
12. Depolymerisation of PET at pilot scale by *Sf* Cut and *Sf* Inv
13. DSC thermograms of PET substrates
14. Thermograms of amorphous film (aFilm) and post-consumer waste (PCW) used in this study
15. Dynamic image analysis of PET powder particle size.
16. HPLC separation trace for the standards TPA, MHET and BHET

#### Supplementary References

**Supplementary Table 1. Accession codes for PETases mentioned in this study.**

| PETase | Accession Code | Ref |
|---|---|---|
| *Bhr*PETase | GBD22443.1 | 1 |
| est119 | BAK48590.1 | 2 |
| *Fs*C | P00590.1 | 3 |
| *Hi*C | HC307279.1 | 4 |
| *Is*PETase | GAP38373.1 | 5 |
| LCC | AEV21261.1 | 6 |
| PE-H | WP_088276085.1 | 7 |
| PET27 | WP_111881932.1 | 8 |
| PHL1 | LT571440.1 ^ | 9 |
| PHL6 | LT571445.1 ^ | 9 |
| PHL7 | LT571446.1 ^ | 9 |
| *Pm*C | Q51718* | 4 |
| *Rg*PETase | WP_085749752.1 | 10 |
| *Sf*Cut (611) | WP_093412886.1 | 11 |
| *Tcur*0390 | WP_012850775.1 | 12 |
| *Tcur1278* | WP_012851645.1 | 12 |
| *Tf*Cut1 | ADV92528.1 | 13 |
| *Tt*CutA | XP_003656017.1 | 14 |

^ ENA database
* UniProt

**Supplementary Table 2. Amino acid sequences for *Sf*Cut and *Sf*Inv.**

| Protein | Amino acid sequence |
|---|---|
| *Sf* Cut | MAEPADVHGPDPTEESITAPRGPFEVDEESVSRLSVSGFGGGTIYYPTDTTDGLFSAVSISPGFTGTQETMAWYG<br>PRLASQGFVVFTIDTITTTDQPDSRARQLQASLDYLVNDSDVKDIIDPARLGVMGHSMGGGGSLKAALDNPALKA<br>AIPLTPWHTTKDFSGVQTPTLIIGAQNDTVAPVSQHAKPFYESLPDDPGKAYLELAGASHLAPNTDNTTIAKFSIA<br>WLKRFLDDDTRYDQFLCPPPENDDSISDYQSTCPYLEHHHHHH |
| *Sf* Inv | MAEPAKVHGPKPTEESITKPRGPFEVNRRSVSRLKVKGFGGGTIYYPTNTTDGLFSAVSISPGFTGTQETMAWYG<br>PRLASQGFVVFTIDTITTTDQPDSRARQLQASLKYLVNKSKVKDIIDPARLGVMGHSMGGGGSLKAALDNPKLKA<br>AIPLTPWHTTKDFRGVRTPTLIIGAQNDTVAPVSQHAEPFYKSLPDKPGKAYLELAGASHLAPNTDNTTIAKFSIA<br>WLKRFLDDDTRYDRFLCPPPRNNKSISDYRSTCPYKEHHHHHH |

**Supplementary Table 3. Extended summary of DSC data with denaturation model fits.** Denaturation data was fitted using Calfitter 2.0[15] to a denaturing model. N -> D means a single irreversible denaturation step, going from native (N) to denatured (D) state without passing through intermediates. $E_{act}$ is the activation energy, $T_{act}$ is the temperature at which the rate of the irreversible step is 1. The 95% confidence intervals, sum of squared residuals (SSR) and degrees of freedom (DOF) statistics are shown.

| | Model | $E_{act}$ (Kcal mol$^{-1}$) | $T_{act}$ (°C) | SSR | DOF |
|---|---|---|---|---|---|
| *Sf* Cut | N -> D | 457.18 ± 0.94 | 76.25 ± 0.01 | 0.06406 | 19236 |
| *Sf* Inverse | N -> D | 378.39 ± 0.35 | 72.64 ± 0.01 | 0.03461 | 20207 |

**Supplementary Table 4. Data collection and refinement statistics (molecular replacement) for X-ray crystallography structure of *Sf* Inv.**

| | *Sf* Inv – 9EWR |
|---|---|
| **Data collection** | |
| Space group | $P2_1$ |
| Cell dimensions | |
| *a*, *b*, *c* (Å) | 46.1, 41.3, 124.1 |
| α, β, γ (°) | 90.0, 99.8, 90.0 |
| Resolution (Å) | 122.27-1.17 (1.32-1.17)* |
| $R_{sym}$ or $R_{merge}$ | 8.1 (76.5) |
| $I / \sigma I$ | 7.2 (1.8) |
| Completeness (%) | 92.5 (62.3)# |
| Redundancy | 6.2 (3.4) |
| **Refinement** | |
| Resolution (Å) | |
| No. reflections | |
| $R_{work}$ / $R_{free}$ | 18.6 / 20.4 |
| No. atoms | |
| Protein | 4031 |
| Ligand/ion | n/a |
| Water | 462 |
| *B*-factors | |
| Protein | 14.6 |
| Ligand/ion | n/a |
| Water | 25.6 |
| R.m.s. deviations | |
| Bond lengths (Å) | 0.0100 |
| Bond angles (°) | 1.97 |

*Values in parentheses are for highest-resolution shell.

#Ellipsoidal completeness

**Supplementary Table 5. Summary of polymer differential scanning calorimetry results.** Melting curves/temperatures were converted to percentage crystallinity by the software, using a reference. Samples were measure in triplicates. Standard deviation shown.

| | Crystallinity (%) |
|---|---|
| Amorphous Film | 7.89 ± 0.47 |
| Amorphous Powder | 9.47 ± 0.73 |
| Crystalline Powder | 32.62 ± 5.16 |
| Post-Consumer Film 1 | 9.64 ± 1.06 |
| Post-Consumer Film 2 | 10.04 ± 0.96 |
| Post-Consumer Film 3 | 8.19 ± 3.92 |

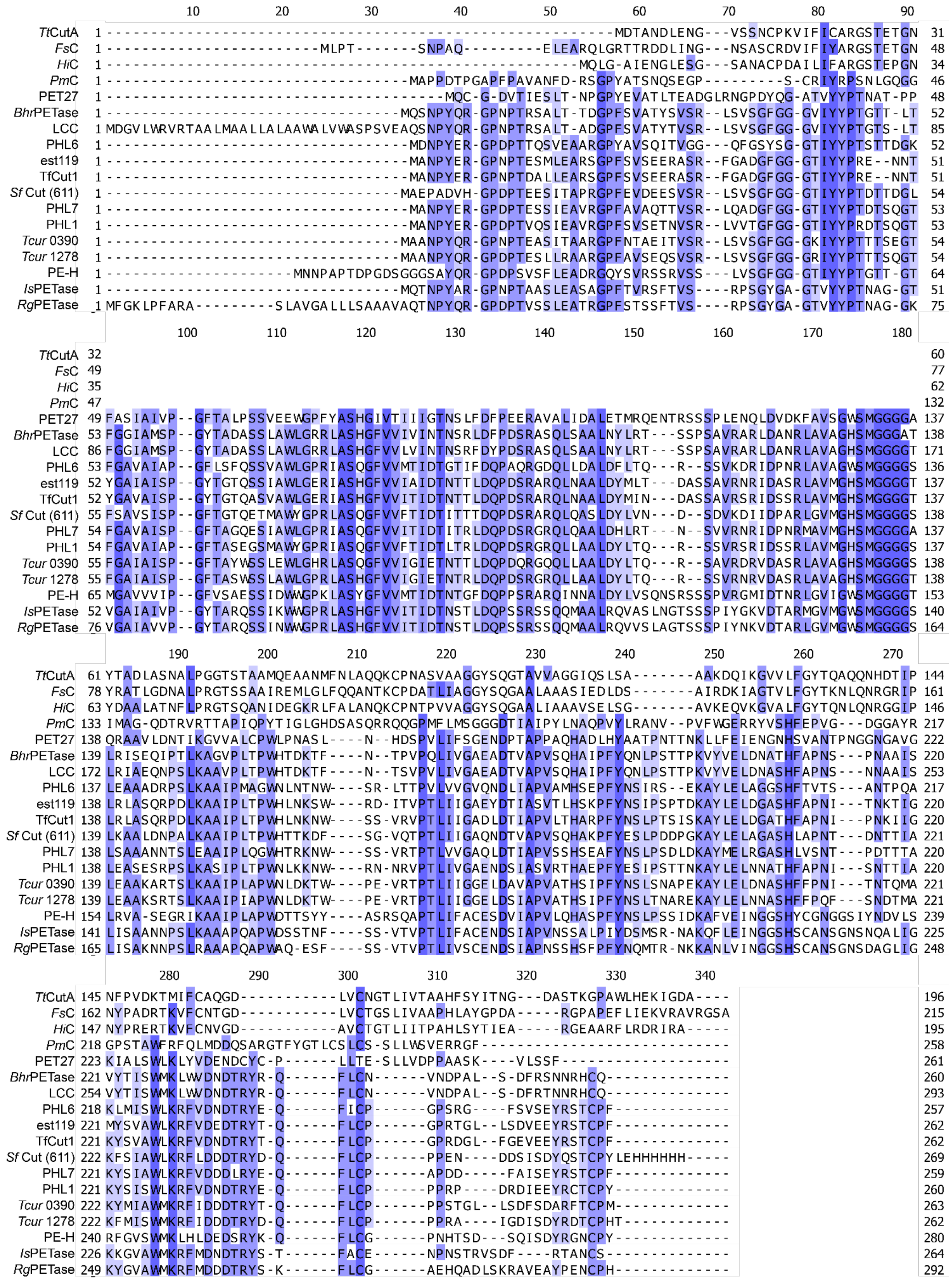


**Supplementary Figure 1. Multiple protein sequence alignment of a set of previously reported PETases.** Alignment is coloured by percentage identity.

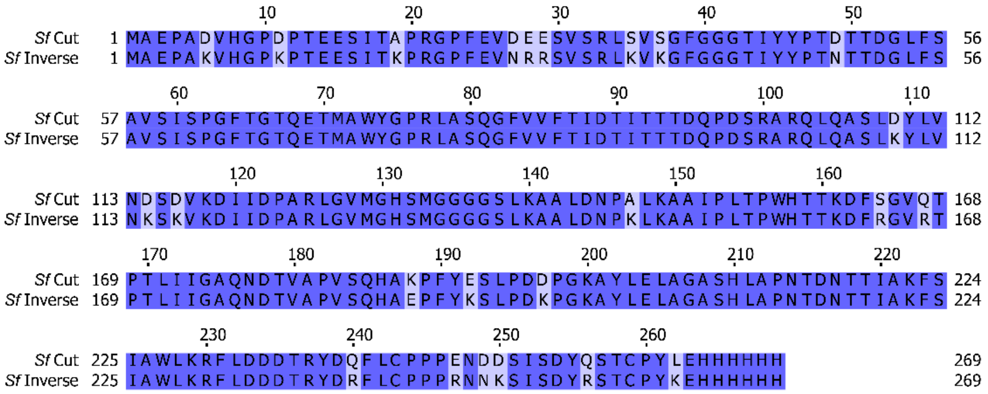


**Supplementary Figure 2. Protein sequence alignment of *Sf* Cut and *Sf* Inv.** Alignment is coloured by percentage identity, with the lighter coloured amino acids highlighting the mutations.

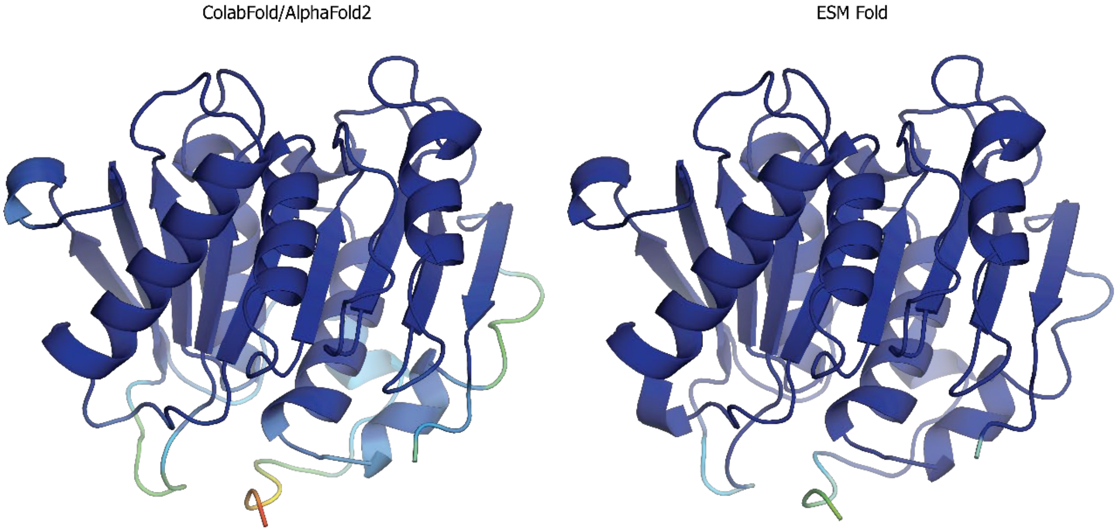


**Supplementary Figure 3. ColabFold[16]/AlphaFold2[17] and ESM Fold[18] predictions of the *Sf* Inv design.** Cartoon representations of the design coloured by pLDDT scores. High confidence regions are shown in dark blue (pLDDT > 0.9), whereas low confidence is indicated in red (pLDDT < 0.5). Overall, both predictions were of high confidence.

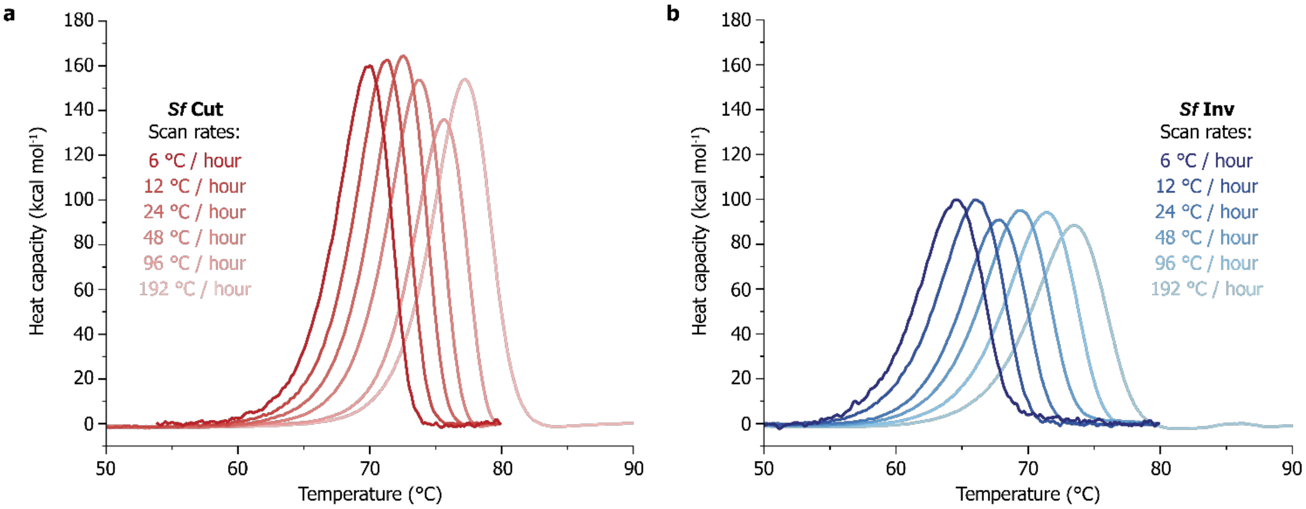


**Supplementary Figure 4. Differential scanning calorimetry curves for *Sf* Cut and *Sf* Inv.** The melt curves for the purified enzymes were scanned at variable rates ranging from 6 °C to 192 °C per hour for both **a** *Sf* Cut and **b** *Sf* Inv.

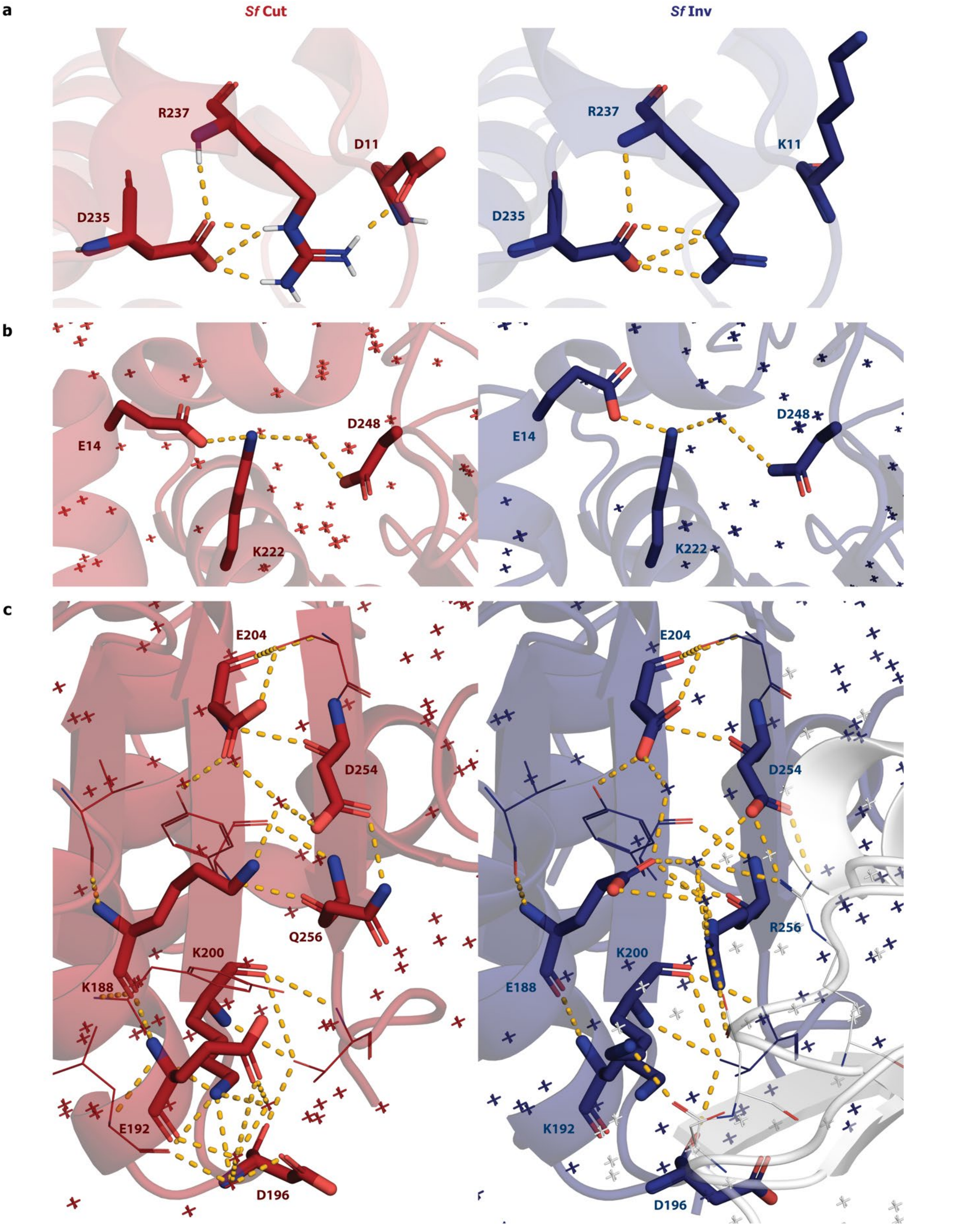


**Supplementary Figure 5. Comparison of polar interactions in modified salt bridges. a** Hydrogen bond between D11 and R237 was lost with the D11K mutation as anticipated in the initial design. **b** D249N mutation is well tolerated and preserved the native contacts. **c** Complex salt-bridge in *Sf* Cut, with multiple interactions happening through coordinated waters (shown as crosses). These are mostly conserved in *Sf* Inv, where the new side chains are also involved in crystal contacts with a symmetry mate (represented in white).

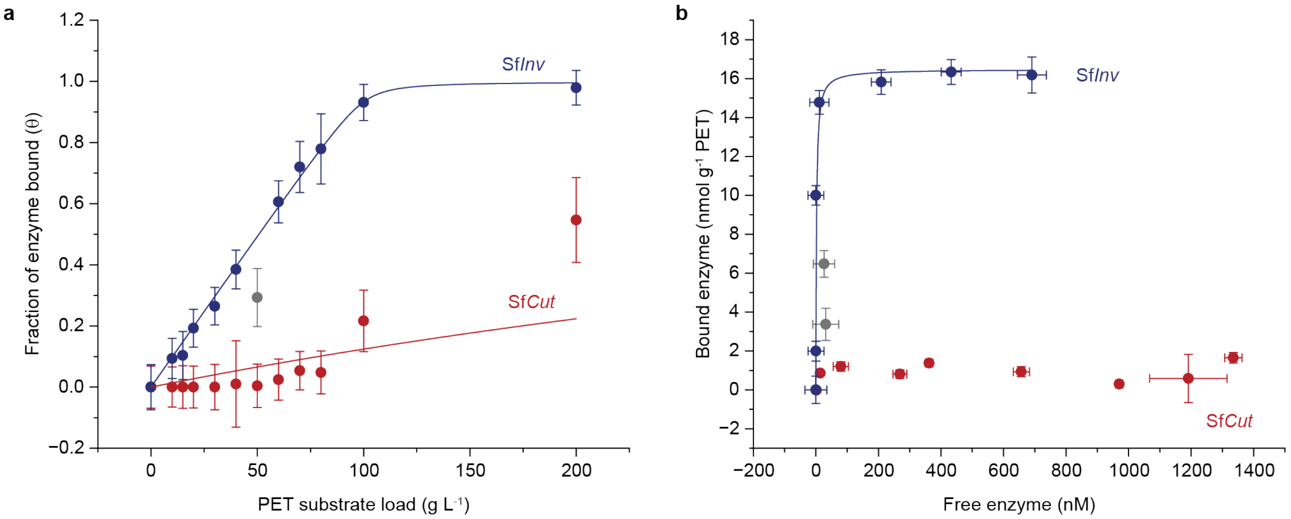


c

| | Enzyme saturation | | | | Substrate saturation | | | |
|---|---|---|---|---|---|---|---|---|
| | $K_d$ (nM) | $\Gamma_{max}$ (nmol $g^{-1}$ PET) | Reduced $\chi^2$ | $R^2$ (COD) | $K_d$ (nM) | $\Gamma_{max}$ (nmol $g^{-1}$ PET) | Reduced $\chi^2$ | $R^2$ (COD) |
| Sf*Cut* | 6128.43 ± 2160.91 [#] | -- | 0.84 | 0.50 | n.d. | n.d. | n.d. | n.d. |
| Sf*Inv* | 4.40 ± 3.64 | 9.94 ± 0.26 | 0.10 | 1.00 | 2.06 ± 3.68 | 16.48 ± 3.49 | 79.89 | 0.46 |

[#]Poor fit quality -- values are for reference only. Fit obtained only when using fixed $\Gamma_{max}$ value of 9.94.

**Supplementary Figure 6. Binding isotherms of *Sf* Inv and *Sf* Cut to amorphous PET powders. a.** Enzymes were incubated at a fixed, saturating concentration (1 μM) with different solids loading of PET powder, and free enzyme measured by the BCA assay. Data is fit to a Langmuir isotherm as previously described[19]. Note: data for *Sf* Cut was insufficient to get reliable metrics from fit, a fixed value of Γmax taken from *Sf* Inv was used to generate curve. It is possible *Sf* Cut demonstrates negative cooperativity in binding to PET powders, but solids loading limitations restrict data collection and analysis **b.** Amorphous PET powder were incubated at fixed, saturating concentrations (50 g/L or 100 g/L for *Sf* Inv and *Sf* Cut, respectively) with different concentrations of enzyme, and free enzyme measured by the BCA assay. Data was fit as described previously[19]. Note: data for bound *Sf* Cut was below the analytical threshold required for analysis (~1 nmol $g^{-1}$ PET). Detailed extracted values from fits are summarised in table below data plots (note poor quality of *Sf* Cut fit statistics).

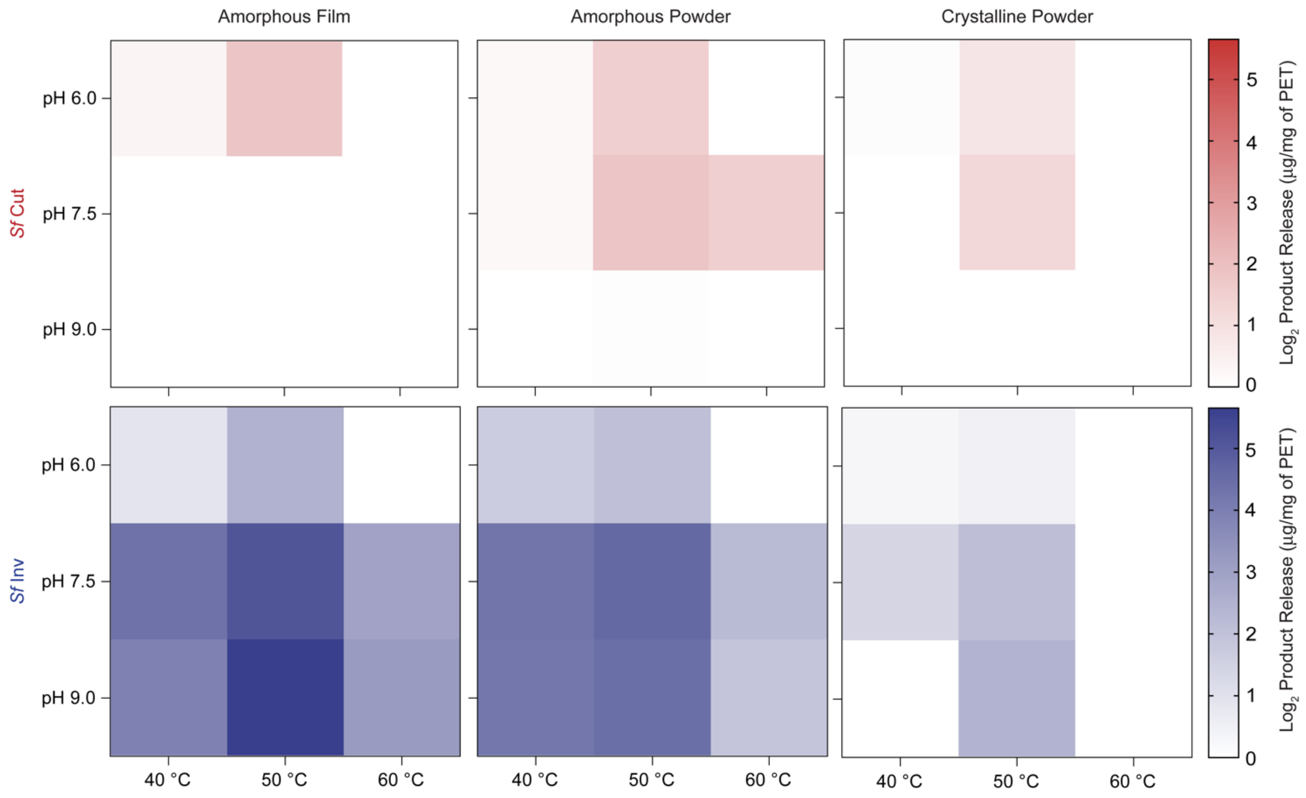


**Supplementary Figure 7**. **Heatmaps of optimum pH and temperature for the enzymes across PET substrates.** *Sf* WT is shown in a gradient of red and *Sf* Inv shown in blue. Reactions were performed at 100 nM enzyme, with 100 mM sodium chloride. For easier comparison activities are shown as $Log_2$ product release (µg/mg of PET)

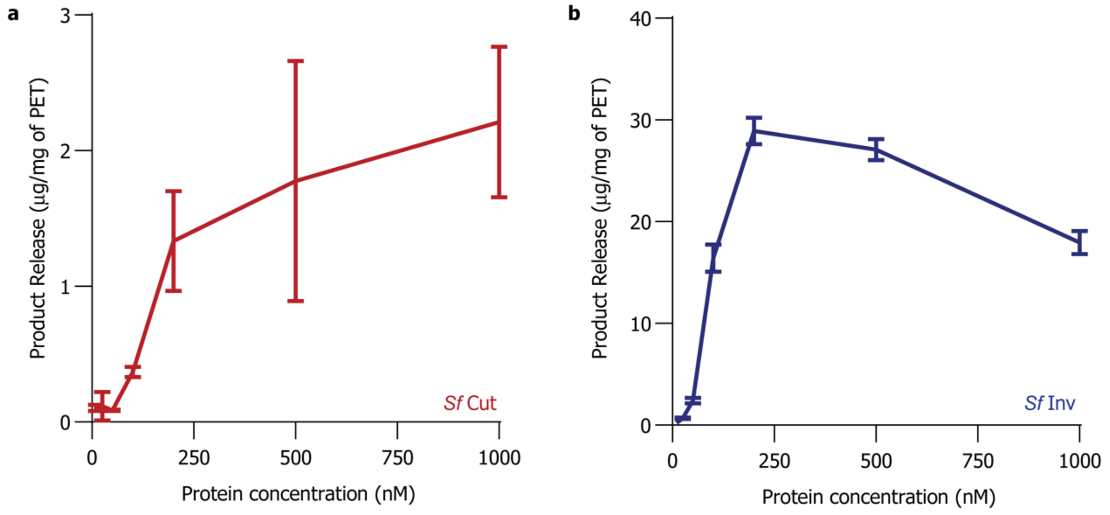


**Supplementary Figure 8. Concentration dependence plots for enzymes on amorphous PET film.** *Sf* Cut (**a**) is shown in red, and *Sf* Inv (**b**) is shown in blue. Measurements were conducted at the enzyme's respective optima on aPET films, at 2.1% solids loading. Both *Sf* Cut and *Sf* Inv saturate above 250 nM, with *Sf* Inv showing evidence of concentration dependent inhibition only at 1µM enzyme.

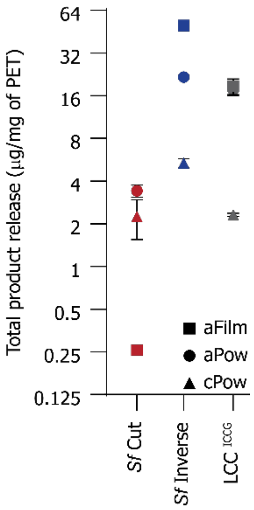


**Supplementary Figure 9. Substrate preference plot for *Sf* Cut, *Sf* Inv and LCC$^{ICCG}$ at 50 °C.** Reactions were performed at 100 nM enzyme and at the optimum pH for the respective enzymes, with 100 mM sodium chloride. Under these conditions *Sf* Inv proves to be more active than LCC$^{ICCG}$[20].

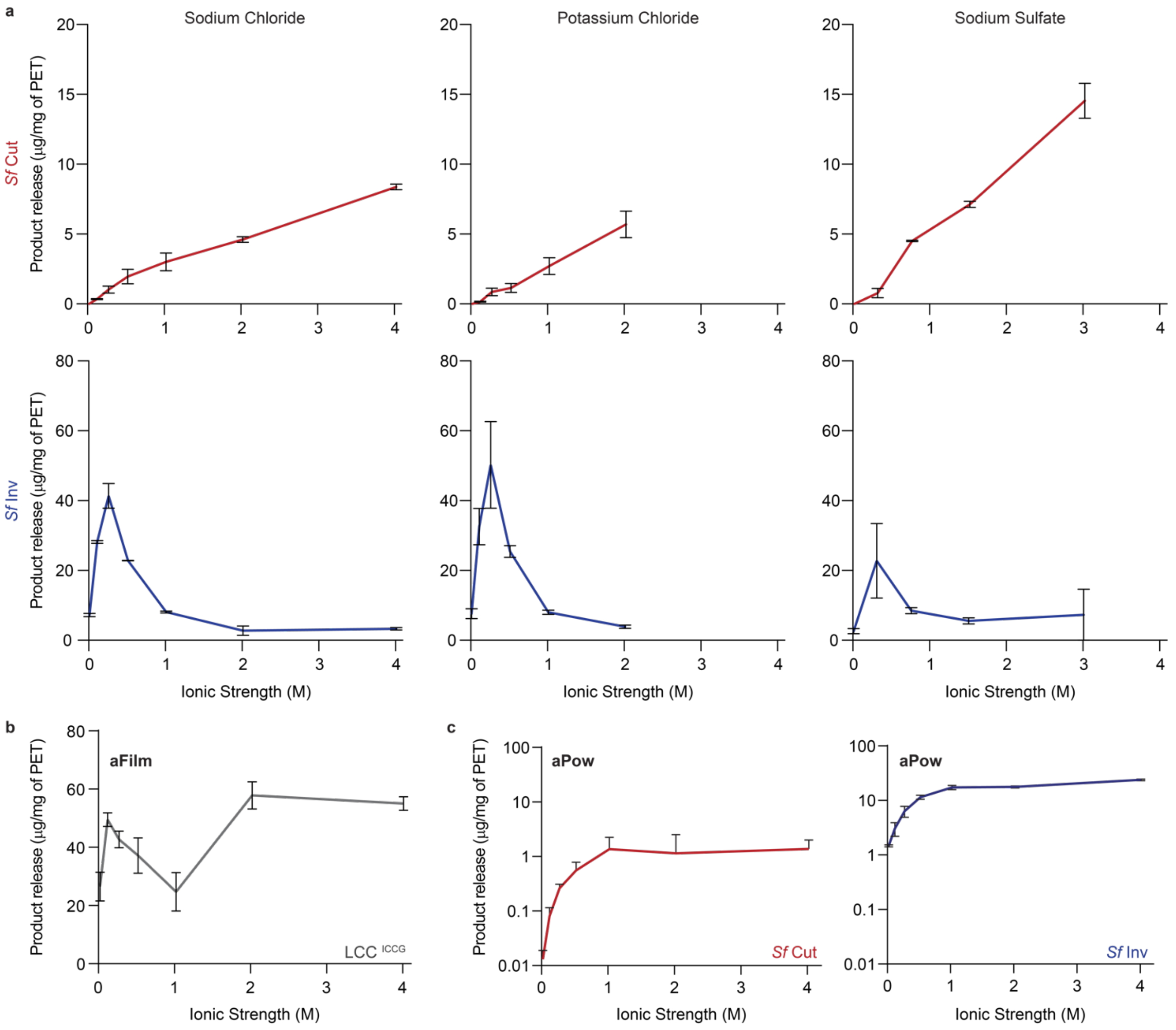


**Supplementary Figure 10. Ionic strength extended – multiple salts and powder comparison. a** Graphs of total product release across different concentration of salts for *Sf* Cut (top three panels) and *Sf* Inv (middle three panels). Three salts were tested (left to right: Sodium chloride, potassium chloride and sodium sulphate). **b** Total product released across different concentrations of sodium chloride for LCC$^{ICCG}$. **c** On amorphous PET films, ionic strength has a positive correlation with total product release for both *Sf* Cut and *Sf* Inv.

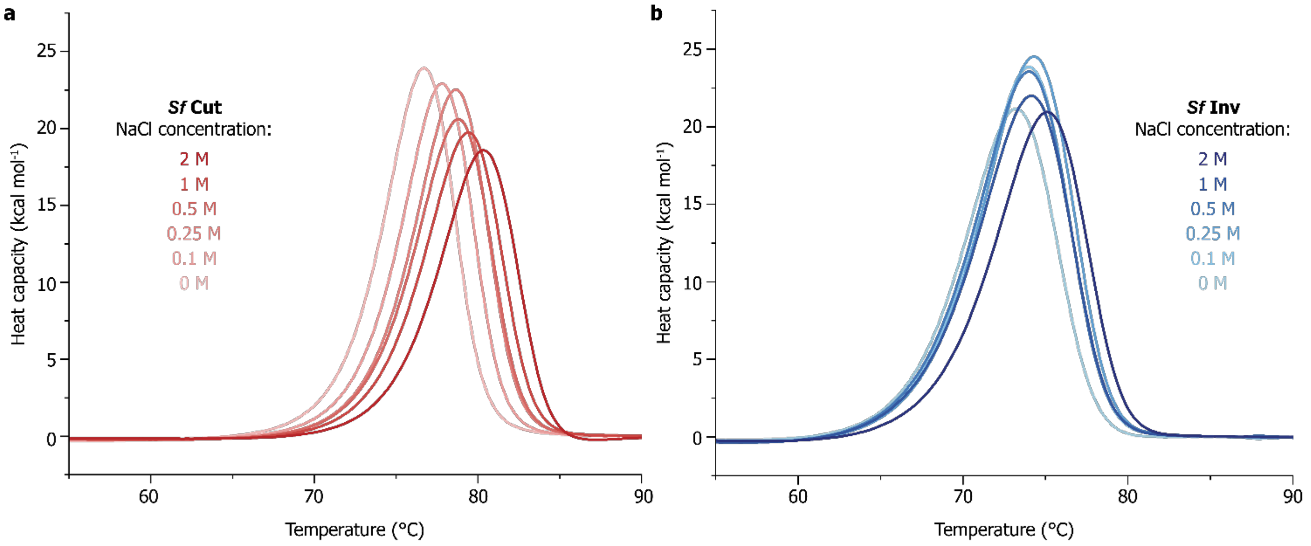


**Supplementary Figure 11. Impact of ionic strength on the apparent $T_m$ of *Sf* Cut and *Sf* Inv.** The purified enzymes were sampled at a single scan rate of 196 C per hour, with different concentration of sodium chloride ranging from 0M to 2M in 50 mM sodium phosphate pH 7.5 buffer. **a** There is a positive correlation between salt concentration and the thermostability of *Sf* Cut, with the apparent $T_m$ increasing from 76.68 °C to 80.34 °C. **b** In contrast, the thermostability of *Sf* Inv is not impacted by increased ionic strength, with the apparent $T_m$ remaining at around 74.12 ± 0.56 °C throughout the titration.

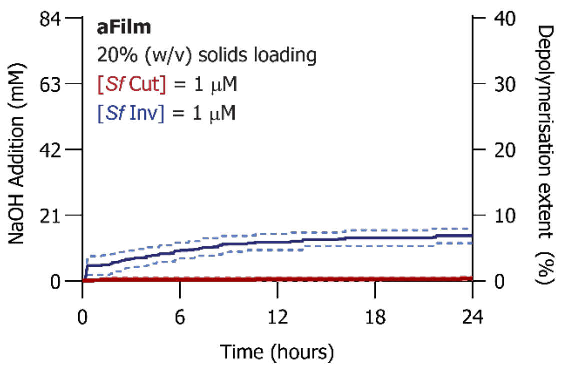


**Supplementary Figure 12. Depolymerisation of PET at pilot scale by *Sf* Cut and *Sf* Inv.** Enzymes were incubated with 20% (w/v) PET substrate, at an enzyme loading of 1 μM final concentration of enzyme (corresponding to around 0.15 mg of enzyme per g of PET). *Sf* Cut is shown in red, whereas *Sf* Inv is shown in blue. Average trace is represented as a solid line, whilst the replicates are represented as dotted lines. *Sf* Inverse was able to achieve the same extent of turnover as it did at a higher enzyme load in Figure 5a, suggesting that lower enzyme loads can be used to achieve equivalent rates of depolymerisation whilst using less resources. *Sf* Cut remained ineffective at tackling plastic films, with less than 1% conversion.

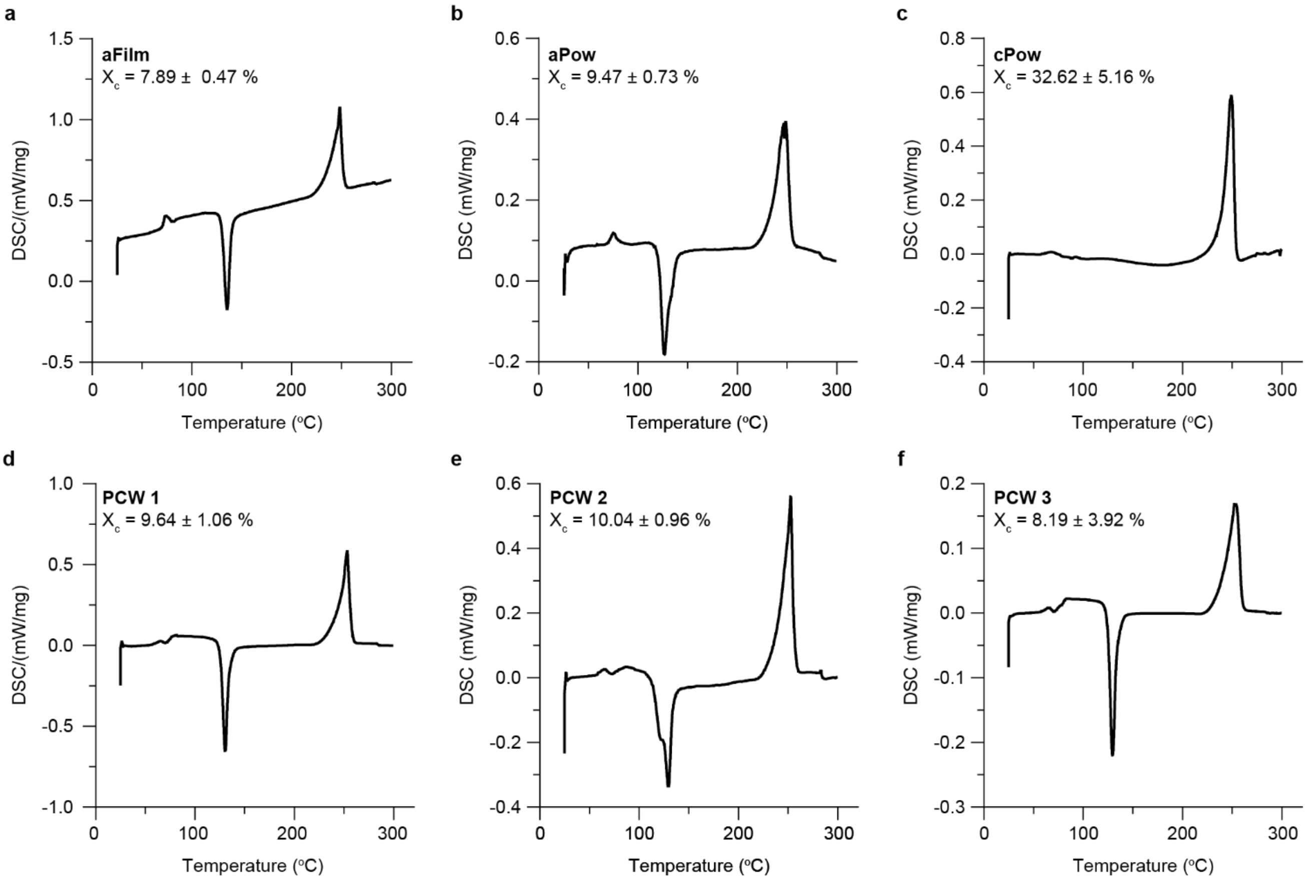


**Supplementary Figure 13. DSC thermograms of PET substrates.** Crystallinity content ($X_c$) for the amorphous film (aFilm, **a**), amorphous powders (aPow, **b**) and semi-crystalline powders (cPow, **c**) substrates used in this study were confirmed by polymer DSC, showing that micronization did not substantially affect the crystallinity of aPow. Post-consumer waste film (PCW, **d-f**) was tested at three different locations on the packaging, where measured crystallinity were within error of each other and comparable to that of the standard aFilm used.

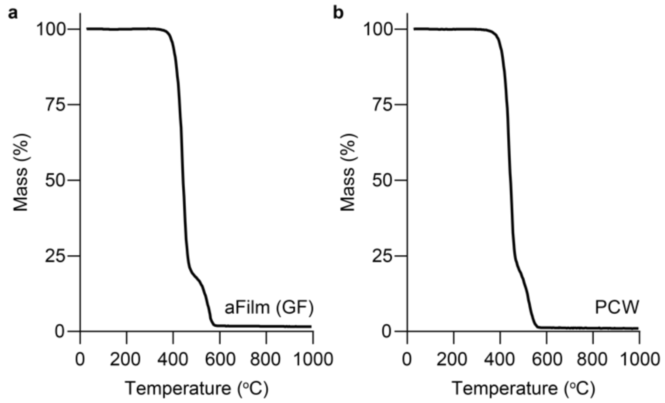


**Supplementary Figure 14. Thermograms of amorphous film (aFilm) and post-consumer waste (PCW) used in this study.** The PCW used matches the onset temperature and mass loss characteristics of the standard aPET film from GoodFellows, confirming that it is PET.

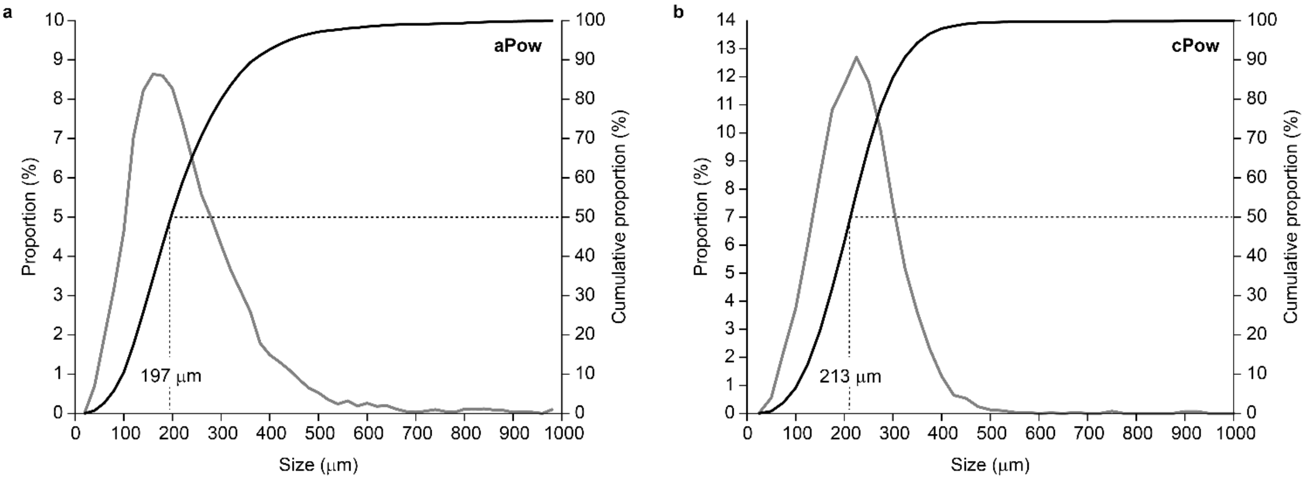


**Supplementary Figure 15. Dynamic image analysis of PET powder particle size.** The cross-sectional diameter population distribution (grey line) and cumulative distribution (black line) for the PET powders used in this study as determined from dynamic image analysis on a Microtrac CAMSIZER X2. The size is the diameter of a circle within the equivalent cross-sectional area as the particle. **a.** Micronized amorphous PET powders and **b.** commercially available semi-crystalline PET powders were measured with similar diameters.

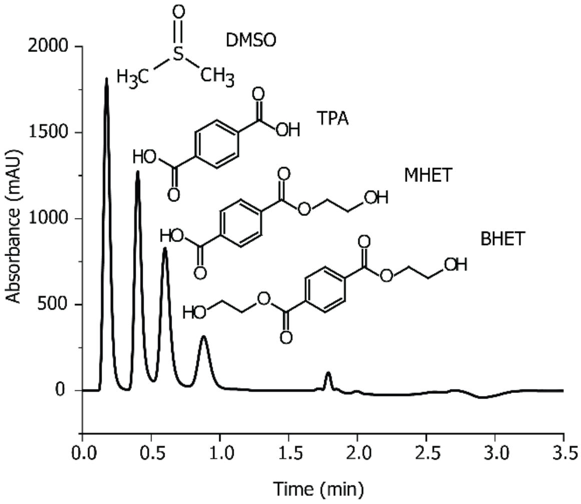


**Supplementary Figure 16. HPLC separation trace for the standards TPA, MHET and BHET.** With this method, TPA eluted after 0.4 minutes, MHET at 0.6 minutes and BHET at 0.9 minutes. The rest of the trace consists of the cleaning and re-equilibration steps. DMSO was used to solubilise the standards.
